# Daily light exposure habits of youth with migraine: A prospective pilot study

**DOI:** 10.1101/2025.04.30.650986

**Authors:** Carlyn Patterson Gentile, Ryan Shah, Blanca Marquez De Prado, Nichelle Raj, Christina L. Szperka, Andrew D. Hershey, Geoffrey K. Aguirre

## Abstract

**Background:** Eighty percent of youth with migraine report photophobia. It is unknown if photophobia leads to light avoidant behavior, and if such behaviors worsen light sensitivity and disrupt sleep. Recently developed wearable, continuous light loggers allow us to address these open questions. We conducted a pilot study to determine the feasibility of measuring light exposure using wearable light loggers in youth with migraine.

**Methods:** Youth 10 – 21 years old with a headache-specialist confirmed ICHD-3 diagnosis of migraine were recruited from CHOP headache clinics. Each participant recorded 7 consecutive days of light logging data from the ActLumus device worn as a pendant around the neck paired with a text-based daily migraine symptom diary during a typical school week between November and March 2024. Validated questionnaires were used to capture any headache and bad headache frequency, headache-related disability, visual sensitivity, fear-of-pain, and sleep disturbance and impairment. Percent time spent within recommended light exposure levels was calculated for the day, 3 hours prior to bedtime, and night. Power analysis was calculated to determine sample size needed for group comparison of baseline characteristics across light intensity and light timing metrics to aid in the design of larger studies.

**Results:** Twenty youth with a median age 17 years [IQR 16, 19], 70% of whom were female completed 7 days of continuous light logger recording and daily headache diary. Data completion rates were high with 136/140 (97.1%) useable days of light logger data, and 100% compliance on the daily headache diary. Participant feedback on the study was positive; 85% would recommend the study to others. On average, participants received recommended light exposure during only 14.5% +/− SD 7.0 of daylight hours. By contrast, participants were more consistently below the recommended maximum light levels 3 hours prior to bed (77.5% +/− 21.6 of the time), and at night (99.1% +/− 2.9 of the time). Youth with chronic migraine (i.e., at least 15 headache days and 8 bad headache days per month) had daily light exposure patterns that were phase shifted 60 minutes later as compared to participants with non-chronic migraine. Power analyses suggest that future tests for differences in light exposure between migraine-characteristic groups (e.g., differing by headache frequency, severity, or disability) will require sample sizes on the order of 50 to 150 to reach 80% power with an alpha of 0.05.

**Conclusion:** Measuring daily light exposure is feasible in pediatric populations with photophobia and reveals intriguing trends in youth with migraine that warrant further study.

## Introduction

Photophobia is a common symptom in neurologic and ophthalmologic disorders including optic neuritis, traumatic brain injury, uveitis, and retinal dystrophy, among others.^1^ In youth with migraine, over 80% report photophobia.^2^ Photophobia may influence the visual diet (i.e. the intensity and timing of light exposure during daily life) as individuals with migraine and photophobia report that they avoid high intensity visual environments.^3^

Variation in visual diet has been associated with health outcome. Exposure to brighter days and darker nights is known to confer reduced mortality risk,^4^ perhaps related to the effect that light has upon circadian biology.^5^ Insufficient daytime melanopic illuminance (short wavelength “blue light”) and evening melanopic illuminance from artificial light both have the potential to disrupt circadian entrainment.^6,7^ Multiple studies have associated nighttime screen use with poorer health-related quality of life and sleep disruption in adolescents.^8–10^ These health effects include higher migraine burden in those who self-report prolonged screen exposure.^11,12^ There has, however, been limited empirical study of how visual diet is altered and potentially influences symptoms in people with migraine and photophobia. It is unknown, for example, if light avoidance behaviors over the long term reduce or enhance photophobia symptoms. Extant studies have the additional limitation that light exposure has been assessed subjectively via surveys or daily diary reports.

Recently developed, wearable light loggers now provide the ability to address these questions using quantified measures of visual diet during daily life.^13–15^ These small, battery-powered devices record the absolute level of light falling upon a detector over a period of many days.

These recordings provide the relative amount of light across wavelengths, supporting inferences regarding the types of light encountered (natural vs. artificial) and the biological effect upon different classes of retinal photoreceptors (melanopsin containing cells vs. cones).

In this study, we measured the visual diet for 20 youth with migraine using a wearable light logger that continuously tracked ambient light exposure over 7 days during a normal school week. We hypothesized that continuous measurement of the visual diet in youth with migraine would be feasible and yield intriguing associations with disease burden in youth with migraine. Specifically, we predicted that a restricted and poorly timed visual diet (i.e., overall less and poorly timed light) would be associated with worse photophobia, chronic migraine, and worse headache-related disability.

## Materials and Methods

### Study Design

This single-center prospective observational pilot study was conducted within the pediatric headache program within the Children’s Hospital of Philadelphia (CHOP) neurology department between October 2024 and March 2025. The protocol received approval from the Institutional Review Board at CHOP.

### Participants

Potentially eligible participants were identified by screening patients being seen in upcoming headache clinics via chart review. Families were contacted via secure phone call three to four weeks before a scheduled clinic visit to determine interest, and if applicable, confirm eligibility and obtain informed consent and assent. Participants were included if they were between the ages 10 to 21 years, ICHD-3 defined migraine with or without aura given by a headache specialist, any headache frequency was allowed. Youth were excluded if they had major neurological conditions besides migraine (e.g. history of epilepsy, stroke, multiple sclerosis), had a recent history of concussion (<3 months), or were starting or weaning off a headache preventive medication (supplement or prescription) or had not been on a stable dose for at least 1 month prior to enrollment.

### Data Collection

After participants received the wearable devices before recording started, a virtual visit was conducted via secure video call. During this visit, study personnel reviewed the proper use of the wearable devices. Participants completed baseline questionnaires through REDCap (Research Electronic Data Capture)^16,17^ hosted by the institution. Instructions for completing the daily headache diary were also provided at this time, then participants started data collection.

The first full day of recording (12 am – 11:59 pm) was considered “Day 1,” and data collection ran for 7 full days. During these 7 days, diary questions were texted between 7pm and 11pm based on participant preference. During data collection, participants continuously wore the light logger and accelerometer while responding to headache diary prompts each evening. The light logger was removed while sleeping to avoid choking or the light logging device being obscured by bedsheets. Participants were instructed to keep the light logger on a bedside table, which is standard for light logger studies,^18^ The light logger was also removed for athletic activities that could be hazardous with a lanyard (e.g., gymnastics), and to avoid water exposure (e.g., showering, swimming) and kept within the same location as the participant during these activities.

After 7 days of data collection, an in person visit coincided with a scheduled headache clinic visit at CHOP. Participants returned the wearable devices, completed any outstanding headache diary entries, and filled out REDCap-based questionnaires. Additionally, the light transmittance of the participant’s personal eyewear was measured, if applicable. Upon completion of the in person visit, participants gave feedback on the study.

### Participant Characteristics

Demographic, medical history, clinical characteristics, and treatment were gathered. Standardized and validated survey data included the CHOP Headache Questionnaire,^19^ including number of any headache days per month and number of bad headache days per month. Light sensitivity was measured using the Visual Light Sensitivity Questionnaire (VLSQ-8), which is an 8-question validated measurement of light sensitivity with possible scores ranging from 8 – 40.^20^ Fear of Pain Questionnaire for Children (FOPQ-C) is a validated metric with was used to measure fear-avoidant responses to pain. After the week of data collection, headache-related disability over the previous month was assessed with the Pediatric Migraine Disability Assessment (PedMIDAS). PedMIDAS has a maximum score of 80 (1-month version), with scores of 3 or less indicating no disability, 3 – 9 indicating mild disability, 10 – 16 indicating moderate disability, and >16 indicating severe disability. PROMIS Sleep Disturbance questionnaire was also filled out to capture the perception of sleep quality and the PROMIS Sleep-Related Impairment questionnaire, which captured symptoms of insufficient sleep (e.g. daytime sleepiness). PROMIS rates no, mild, moderate, and severe symptoms based on T-scores.^21^ Data were also collected on chronotype, using the standardized chronotype questionnaire.^22^

### Headache Diary

Daily migraine symptoms, acute medication use, and function were recorded with a validated text-based Daily Headache Diary^23^ with additional questions to capture device wear compliance, and light-blocking lens use. Daily headache diaries were filled out via text message using the HIPAA-compliant Twilio platform hosted at CHOP. Specific questions included “Have you had a headache today” (yes or no), “Has your headache gotten in the way of your school, home, or social life today” (yes or no), and “Rate your light sensitivity (0 – 5).”

### Light Logger Data collection

ActLumus devices (Condor Instruments, São Paolo, Brazil), which are wearable light loggers that provides 24-hour continuous collection of photopic illuminance and melanopic equivalent daylight illuminance (mEDI).

#### Device wear and data collection

ActLumus devices (Condor Instruments, São Paolo, Brazil), which are wearable light loggers that provide 24-hour continuous collection of photopic illuminance and mEDI. ActLumus devices were worn as a pendant around the neck, providing more ecologically valid measurements that are not obscured by shirt sleeves and are closer to the visual plane, which is critical for capturing non-image forming effects of light.^14,24^ Data were collected at a sampling rate of 1/minute to maximize temporal resolution while still providing for 7 days of continuous recording.^15^

#### Data processing

Data were visually inspected and excluded if participants reported not wearing and being away from the Actlumus device for more than 2 hours in a day. ActLumus data were pre-processed using ActiLab Software (Condor Instruments, São Paolo, Brazil). Specifically, melanopic and photopic lux values were derived from 10 light sensing channels that detect different wavelengths of light for every minute of data capture. Then, melanopic and photopic illuminance was processed using custom software developed in Matlab (Mathworks). Log transformation was performed to accommodate large shifts in illuminance levels.^13^

#### Light metrics

Two features of the visual diet were considered: 1) the intensity of light exposure, 2) the timing of light exposure. Light metrics were chosen that summarize continuous light exposure data to capture these two features of the visual diet.^13,25^

Light intensity was represented by photopic luminous exposure and time spent exposed to outdoor light levels. Photopic luminous exposure captures the total 24-hour photopic light exposure (kilolux*hr), which provides a measurement of overall light exposure. Time spent in outdoor light was defined as time spent in photopic illuminance >1,000 lux because outdoor light ranges from 1,000 lux on a cloudy day to 100,000 lux on a bright sunny day, while indoor light is generally below 500 lux.^25–27^

Light timing was defined by percent time spent within recommended mEDI limits. A minimum 250 melanopic lux is recommended during daytime hours, while a maximum of 10 lux in the evening 3 starting three hours before bedtime, and 1 lux or less at night is recommended to support optimal timed melatonin release.^28^ These recommendations were based on expert-scientific consensus and supported by the sensitivity of human “non-visual” responses to ocular light. In this study, we chose fixed definitions of “day” defined as 7 am – 5 pm, “pre-bedtime” defined as 8 pm – 11 pm, and “night” defined as 12 am – 6 am, based on the diurnal motion of the sun and the structured schedule imposed by school. Gaps in time were left between “day,” “pre-bed,” and “night” definitions to allow for some variability across individual schedules.

Based on differences observed in participants with and without chronic migraine (see *Results*), we added mean mEDI for transition periods including around the time of first morning light exposure or “lights on” (6 am – 8 am), the afternoon typically around the end of school (3 pm – 6 pm), and around “lights out” (9 pm – 12 am).

### Validation of Light Logger measurements

We validated the tabular spectral sensitivity functions of the ActLumus device. To do so, we measured a standard light source using both the ActLumus and a calibrated spectrophotometer (SpectraScan® PR-670, JADAK, North Syracuse, NY). The light source was the output of an 8-channel, digital light synthesizer (CombiLED, Prizmatix, Tel Aviv, Israel) delivered via liquid light guide into a light integrating sphere (LabSphere, North Sutton, NH). The spectral radiance of the light source was measured at 2 nm resolution using the PR-670. Combining this spectrum with the ActLumus tabular sensitivity functions (provided by the manufacturer), and converting from radiance to irradiance, provided a model prediction of the ActLumus sensor counts. We then measured the light source using the ActLumus, and compared the obtained and predicted sensor counts. We found the ActLumus counts to be in excellent agreement with the prediction (Pearson’s R = 0.9982). We were, however, limited to validating 9 of the 10 ActLumus channels, as the PR-670 does not measure the infra-red sampling range of the 10^th^ channel.

### Statistical Analysis

No *a priori* sample size calculations were performed as this was a pilot study to determine sample size for future studies. Descriptive statistics for continuous variables included median with interquartile range for non-normal continuous distributions and mean with standard deviation for continuous variables with normal distribution. Proportions were reported for categorical variables. Continuous light logger data was graphically represented as the mean with 95% confidence intervals (CI) determined by bootstrap analysis. Shifts in the temporal profile between weekdays and weekends and between youth with and without chronic migraine were estimated by determining the correlation between a shifted version (+/− 100 minutes) of the first group and the second group and taking the highest correlation value (which was r > 0.98 for both comparisons).

Power analysis of migraine characteristics included comparison between the following groups using two-sample two-tailed t-test: (1) those with at least 15 headache days per month and 8 migraine days per month based on ICHD-3 criteria for chronic migraine^29^ compared to those who did not meet this criteria, (2) those with no-to-mild versus moderate-to-severe headache-related disability as defined by PedMIDAS, (3) those with low visual sensitivity (VLSQ-8 ≤ 24) versus high visual sensitivity (VLSQ-8 > 24), (4) those with no or mild versus moderate-to-severe sleep disturbance defined by PROMIS, (5) those with no or mild versus moderate-to-severe sleep impairment defined by PROMIS. Power analysis was calculated by resampling the pilot participants within each group with replacement 1,000 times for sample sizes ranging from 5 to 75 per group. The sample size needed to achieve 80% power with an alpha of 0.05 was determined.

## Results

### Participant characteristics

Twenty youth with migraine participated in this study. Demographics and clinical characteristics were reported (Table 1). Participants were a median age of 17 years old [IQR 16, 19] and were 70% female, and reported a median of 17 [IQR 6, 30] any headache days per month, and 5 [IQR 2, 15] bad headache days for the previous month at the start of recording. Headache-related disability (measured by 1-month PedMIDAS scores) had a median score of 16 [IQR 8, 29], which is consistent with moderate disability.^30,31^ Median fear-of-pain (FOPQ-C) score was 39 [31, 54], placing most youth in the moderate-to-severe range, consistent with chronic pain conditions, including migraine.^32^ One participant (5%) reported frequent use of acute medications placing them at risk for medication adaptation headache. All participants were on at least one, and many were on a combination of pharmacologic agents for headache prevention. Supplements, OnabotulinumtoxinA, SNRIs/SSRIs, and CGRP blocking medications were the most common.

**Table 1.**
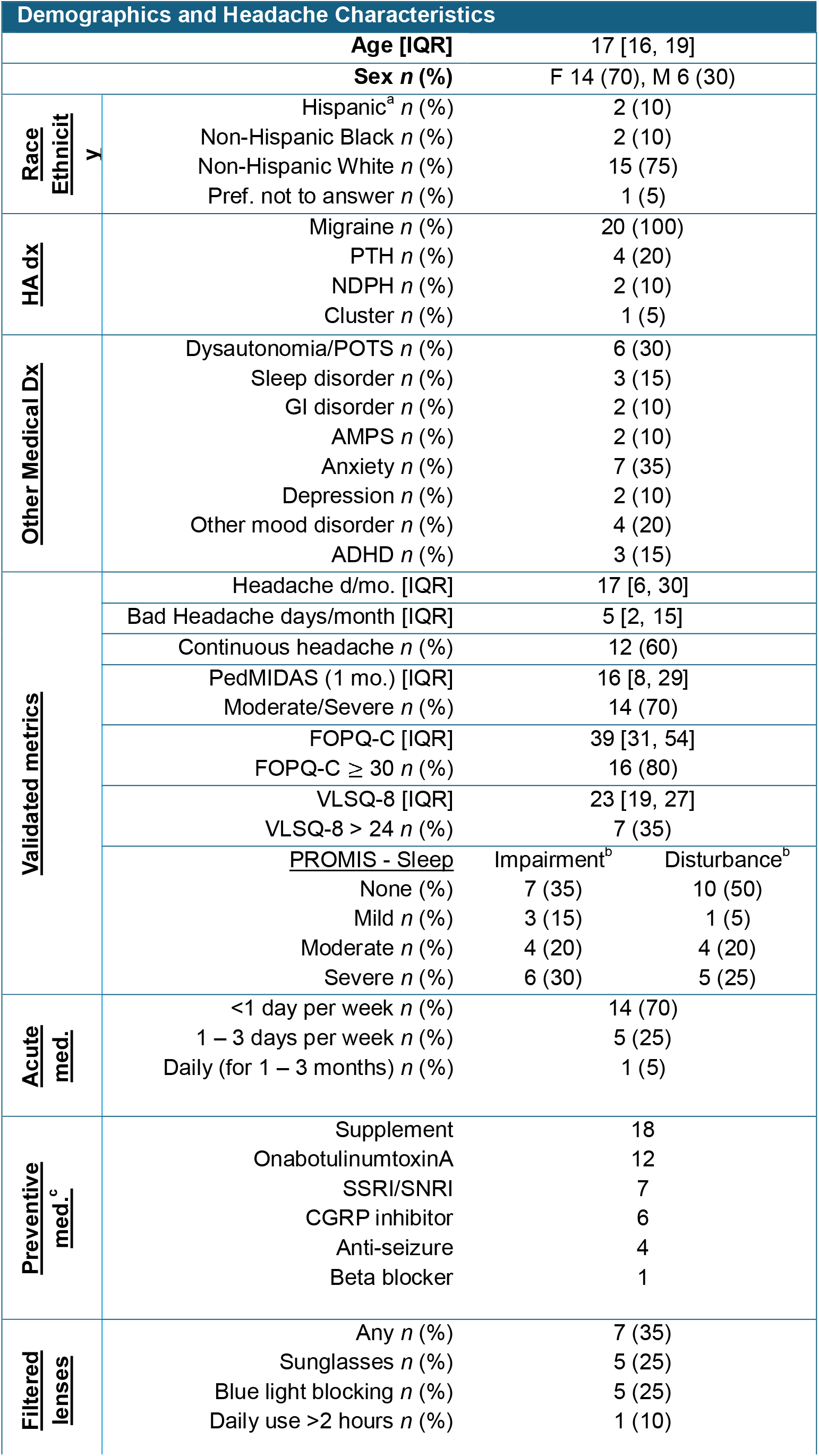
Participant demographics, baseline headache characteristics, and treatment. ^a^For the two participants who indicated they were Hispanic, both selected “prefer not to answer” for race. ^B^Sleep impairment measures symptoms of poor sleep including fatigue and daytime sleepiness, while sleep disturbance measures difficulties falling and staying asleep. ^c^Many participants were on more than one preventive therapy. CGRP included fremanezumab (3), galcanezumab (1), atogepant (2), Anti-seizure included topiramate (4) and lamotrigine (1), supplements included Magnesium (7), Riboflavin (3), B-complex (2), Vitamin D (2), Vitassium (2), Co Enzyme Q10 (1), Melatonin (1), beta blocker was propranolol (1). PTH = post-traumatic headache; HA = headache; Dx = diagnosis; med. = medication; NDPH = new daily persistent headache; POTS = Postural Orthostatic Tachycardia Syndrome; AMPS = Amplified musculoskeletal pain syndrome; ADHD = attention deficit hyperactivity disorder; CGRP = Calcitonin gene-related peptide; SSRI = serotonin reuptake inhibitor; SNRI = serotonin and norepinephrine reuptake inhibitor.

**Table 2.** Headache Diary Data. Includes overall counts for symptoms across all participants and individual participant data. Five participants reported light blocking glasses use, but only one used light blocking glasses throughout the day every day, with the remaining 4 using on an as needed basis. Med. = median; IQR = interquartile range.

| Headache Diary Metrics Collected Over 7 Days |  |  |
| --- | --- | --- |
| Total n (%) | Days recorded | 140 |
|  | Headache days | 98 (70.0) |
|  | Bad Headache days | 63 (45.0) |
|  | Days an acute med was used | 30 (21.4) |
|  | Days with HA-related disability | 42 (30.0) |
| | Days with light sensitivity score $\geq 3$ | 69 (49.2) |
| Participant Med. [IQR] | Headache days/week | 6 [3, 7] |
|  | Bad Headache days/week | 3 [1, 6] |
|  | Days with HA-related disability | 2 [1, 4] |
|  | Light Sensitivity Score (0 – 5) | 2 [1, 3] |
|  | Pain Score (0 – 10) | 4 [0, 6] |

### Daily headache diary data

All 20 participants had 100% compliance with headache diary prompts. Across all 140 recording days, 70.0% were headache days, 45.0% were bad headache days, and acute medications were used on 21.4% of days. Thirty percent of the days were associated with headache-related disability, and nearly half (49.2%) were associated with at least moderate light sensitivity. Youth reported a median of 6 headache days [IQR 3, 7], and 3 bad headache days [IQR 1, 6], indicating they had high any headache and bad headache frequency during recording week, consistent with baseline measurements. Median daily light sensitivity score was 2 [IQR 1, 3] indicating mild light sensitivity, and median pain score was 4 [0, 6] indicating moderate pain.

Seven participants reported using light blocking lenses. Of those, only one participant used light filtering glasses continuously, with the remainder using glasses a maximum of 1-2 hours a day on an as needed basis. There were challenges with lens transmittance measurements, so these values could not be reported.

### Light logger measurements

Each participant completed 7 days of recording for a total of 140 days. For one participant, 4 days of recording were excluded because of being away from the device for more than 2 hours, which was consistent with visual inspection of the data (3 days) and 1 additional day based on visual inspection where the device did not record (1 day). This left 136/140 (97.1%) of days with usable ActLumus data across 20 participants.

### Intensity of light exposure

Two metrics of photopic illuminance were used to characterize the overall light intensity (Figure 2). Total photopic luminous exposure was calculated, which represents the integrated exposure to photopic light in a 24-hour period. The mean total luminous exposure across participants was 6.2 klux*hr, with a wide range between 0.2 and 16.9 klux*hr. Time youth spent in outdoor light was also estimated, which was defined as light exposure of 1,000 lux photopic illuminance or greater, as indoor light is typically below 500 lux. This provided an intuitive measure of periods spent in bright light. The mean total time spend in outdoor light across participants was 42 minutes per day [range 0 to 108 minutes].

**Figure 1.**
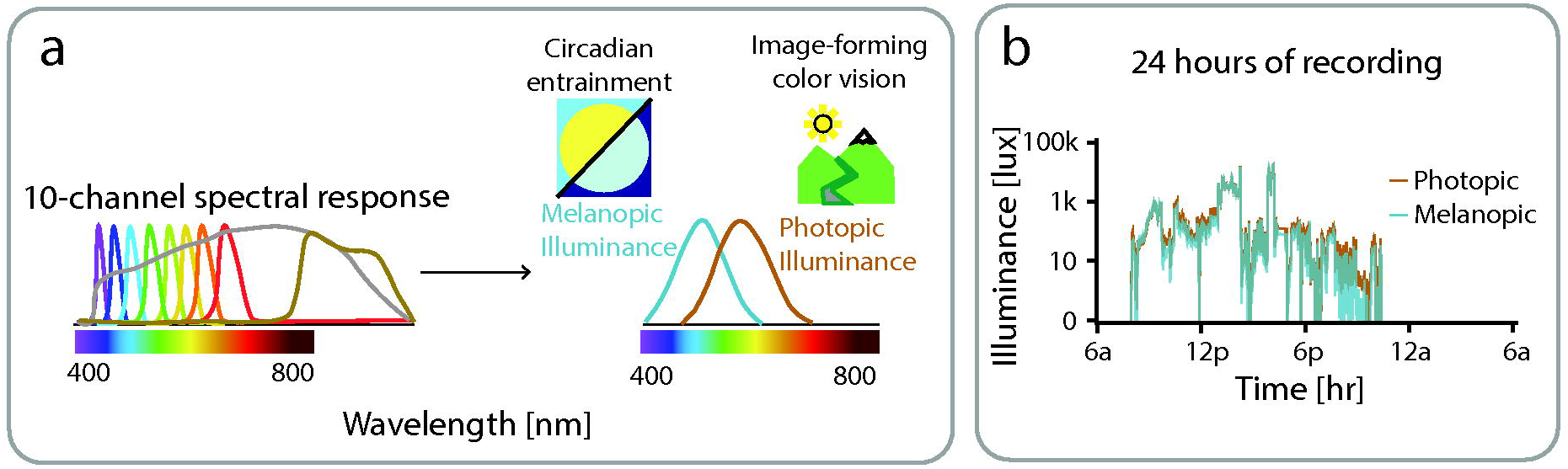
Actlumus light recording. (a) Illustration of the Actlumus light sensor consisting of 10 Channels with different spectral sensitivities that sample at a rate of 1/minute. These measurements are differentially combined to derive melanopic illuminance that is important in circadian rhythm signaling and photopic illuminance that is the basis for image forming daylight color vision. (b) An example participant of continuously recorded of photopic and melanopic illuminance over a 24-hour recording period. Light exposure increases to measurable levels (:2: 0.01 lux) starting at 7am and then drops below measurable levels at 11pm that follows a typical 24-hour sleep/wake cycle. Under most circumstances, photopic and melanopic illuminance are highly correlated (e.g. daylight). However, a dissociation can be seen in the evening, which may reflect use of screen filters with phone use or certain forms of artificial light.

**Figure 2.**
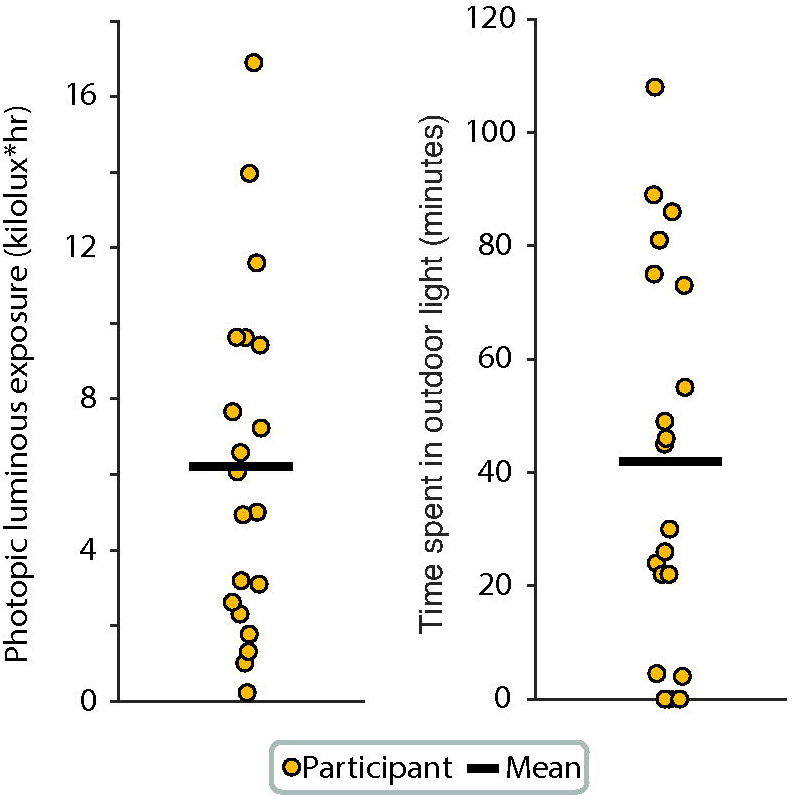
Light intensity. Photopic illuminance was used to calculate light intensity metrics as these have been used as the standard for designing lighting spaces. Each participant represents the median photophic luminous exposure (left panel) and the median time spent in outdoor light levels (right panel) across the 7-day period for each participant. The mean (black line) is also shown. Median was used for each participant due to the non-normal distribution of the data across 7 days, and mean was used for between participants because values approximated a normal distribution.

### Timing of light exposure

Different levels of light exposure are recommended during the day, three hours before bed, and night: a minimum light exposure of 250 melanopic lux during daytime hours; a maximum of 10 lux starting three hours before bedtime; and 1 lux or less at night.^28^ We therefore calculated the proportion of time within these recommended limits during daytime (7a – 5p), pre-bedtime (8a – 11p), and nighttime (12a – 6a) hours. Timing was selected based on typical school schedules, the diurnal pattern of the sun at the location of the study. These definitions were supported by the timing of light exposure observed across participants within a 24-hour period (Figure 3a).

**Figure 3.**
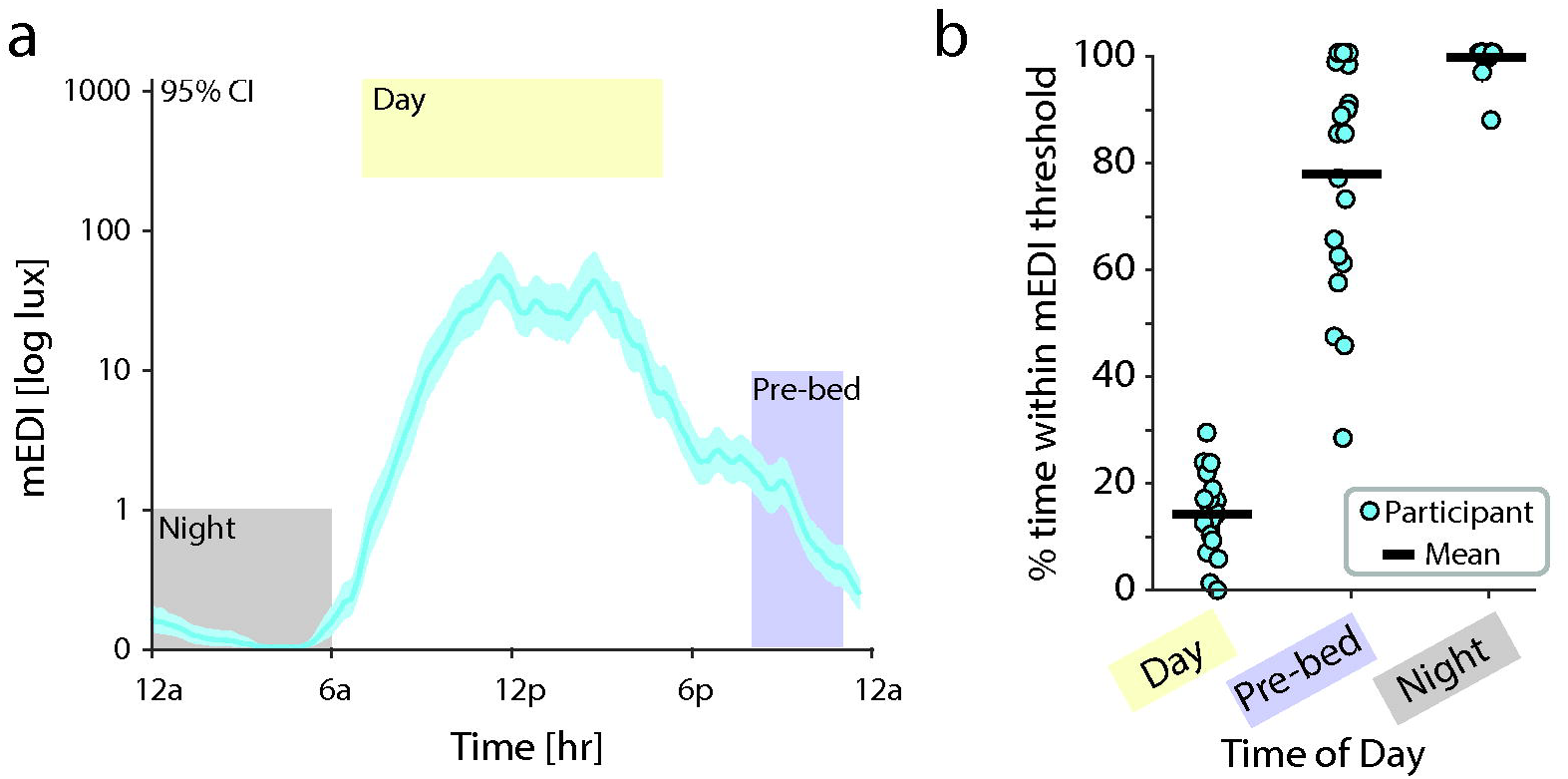
Light timing. Melanopic illuminance (light blue) was used to measure the timing of light exposure because this is important for circadian entrainment. (a) Mean mEDI of 20 participants averaged across 7 days of recording. mEDI was converted to log scale, then the mean was taken for each participant across the 7 days of recording, then the mean across participants calculated for every minute using a sliding window (width 30 minutes). Error bars represent 95% Cl between participants. Yellow (day), blue (pre-bed),and gray (night) squares indicate recommended mEDI levels based on time of day. (b) Percent time spent within recommended mEDI based on time of day: above 250 lux mEDI during the day (left), below 10 lux starting 3 hours prior to bed (middle), and below 1 lux at night during sleep (right). Mean across participants (black) and individual participants averaged across 7 days (light blue circle) are shown. mEDI = melanopic equivalent daylight illuminance.

Participants spent an average of 14.5% +/− SD 7.0% of daytime exposed to the recommended minimum mEDI of 250 lux (Figure 3b). Percent time spent within recommended levels improved substantially in the pre-bedtime and night hours, with youth spending an average of 77.5% +/− SD 21.6% of the time the maximum recommended mEDI of light pre-bedtime, and 99.1% +/− SD 2.9% of the time during night hours.

We next compared melanopic lux patterns and reported sleep and wake times across a 24-hour period to assess the relationship between light exposure and sleep. Weekday versus weekend light exposure habits were compared as patterns are expected to differ given the structure imposed by school on the weekdays but not weekends. As expected, light exposure levels were shifted by an average of 49 minutes later in the day on the weekends compared to the weekdays (Figure 4a). This shift was consistent with later bedtimes, sleep times, and wake-up times reported in the Chronotype questionnaire (Figure 4b).

**Figure 4.**
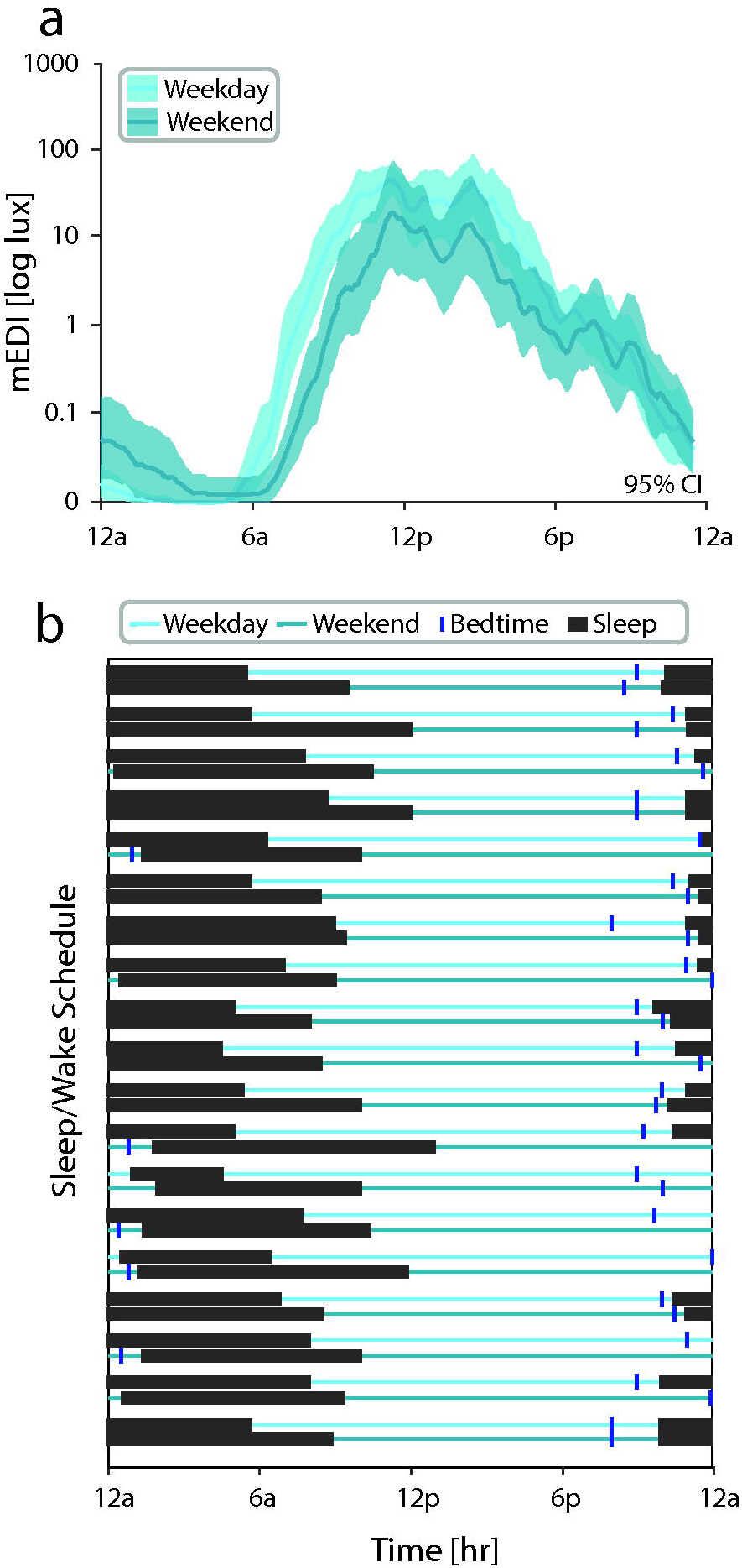
Comparison of light exposure and sleep/wake schedules. (a) Comparison of **mEDI** during weekdays and weekends. (b) Sleep/wake habits for each participant on weekdays (light teal, upper) and weekends (dark teal, lower). Reported bedtime (dark blue vertical line) and sleep period (black box) are shown. **mEDI** = melanopic equivalent daylight illuminance.

To determine if there were any emerging differences in youth with migraine based on headache frequency, we compared the temporal profiles of melanopic illuminance of youth with chronic migraine (15 or more headache days per month with 8 or more bad headache days per month; *n* = 8) to youth who did not meet these criteria (*n* = 12; Figure 5). Youth with chronic migraine demonstrated a shift in the temporal profile of melanopic illuminance by 60 minutes later on average, and an increase in the intensity of melanopic illuminance compared to those without chronic migraine. This shift led to reduced exposure to melanopic illuminance in the early morning (6 am – 8 am) and an increased exposure to melanopic illuminance in the afternoon and evening for youth with chronic migraine, based on non-overlapping 95% confidence intervals as determined by bootstrap analysis. We did not pursue additional statistical testing as the goal of this study was to determine sample sizes needed to appropriately power future research.

**Figure 5.**
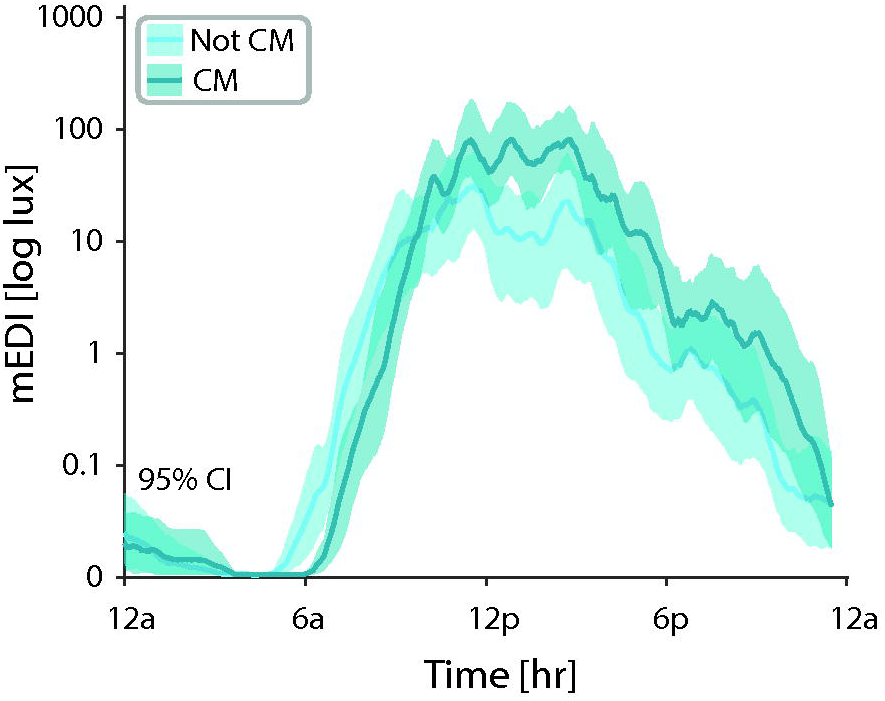
Differences in mEDI exposure in low frequency and high frequency (i.e. chronic) migraine. Comparison of mEDI across participants with CM (dark teal) compared to those who did not meet criteria for CM (light teal). mEDI for each minute was averaged using a sliding window (width 30 minutes). Error bars represent 95% Cl by bootstrap analysis. CM = chronic migraine; mEDI = melanopic equivalent daylight iiluminance.

**Figure 6.**
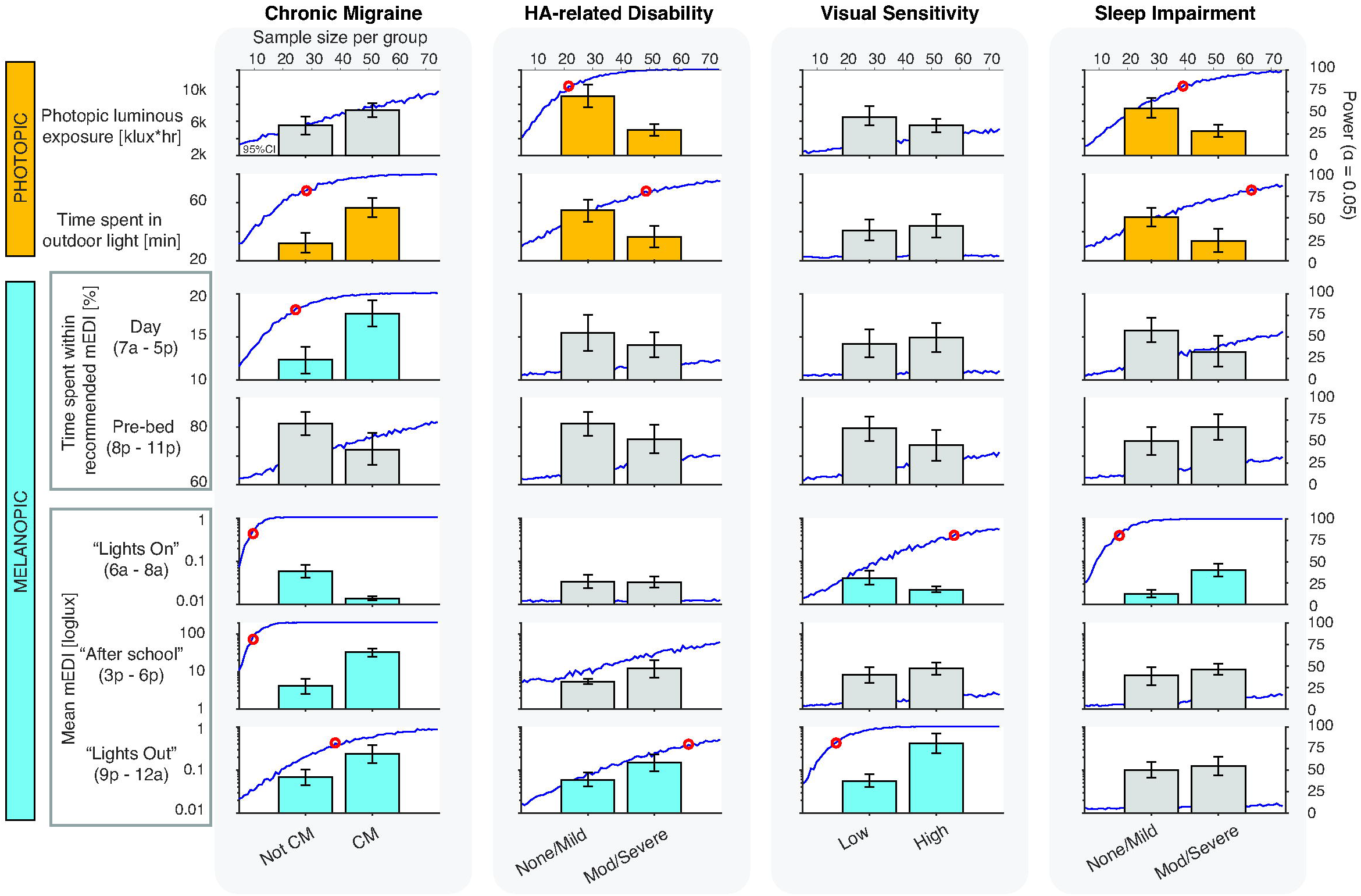
Power analysis of group comparison of clinical characteristics across light exposure metrics. Bar graphs represent mean and 95% Cl for simulated light metric data with 75 participants per group. Dark blue line represents the power for an alpha of 0.05 for sample sizes ranging from 5 to 75 per group. Red circles represent the sample size where 80% power is achieved. Metrics that reached 80% power within 75 participants per group are shown in orange (photopic metrics) and light blue (melanopic metrics) and the minimum sample size that achieves this power threshold is reported on the figure. Metrics that did not achieve this within 75 participants are shown in gray. CM = chronic migraine; Mod = moderate. **mEDI** = melanopic equivalent daylight iiluminance.

### Power analysis

Light exposure may offer a modifiable risk factor for increased disease burden in youth with migraine that could be a target for future interventions. Therefore, it is important to understand if and how light exposure differs with disease burden in youth with migraine. As a first step, we conducted analyses to determine sample sizes needed to adequately power such comparisons.

We compared the following groups: youth with versus without chronic migraine, low versus high headache-related disability, low versus high visual sensitivity, low versus high fear-of-pain, low versus high sleep disturbance, and low versus high sleep impairment. We compared simulated measurements of total photopic illuminance, time spent in outdoor light, percent time within recommended daytime and pre-bedtime light levels across these groups. Percent time spent within recommended nighttime light levels was removed because all but 2 participants were within the recommended range >99% of the time indicating very little variability in this metric.

Based on the pattern of mEDI exposure throughout the day, mean mEDI was also calculated during periods of transition, including 6 am – 8 am (“lights on” or the time frame of the start of morning light exposure), 3 pm – 6 pm (period after school), and 9 pm – 12 am (“lights out” or time frame of the onset of nighttime light levels). Power analysis was conducted by simulating data for sample sizes ranging from 5 to 75 participants per group by resampling from the original 20 participants with replacement.

Group comparison of youth with and without chronic migraine, with low versus high headache-related disability, low versus high visual sensitivity, and low versus high sleep impairment had at least two differences light metrics that reached 80% power with 75 participants or less. Their simulated light exposure profiles are shown (Figure 5). Percent time spent within recommended light for 3 hours prior to bedtime had sufficient power to identify differences between youth with low versus high sleep disturbance (*n* = 30 to achieve 80% power); youth with sleep disturbance spent more time above recommended mEDI. No light metrics had sufficient power to identify differences between youth with low compared to high fear-of-pain scores with 75 participants per group or less.

### Participant Feedback

Seventeen (85%) participants agreed or strongly agreed with the statement “I would recommend somebody to participate in this study,” while 3 (15%) strongly disagreed. Specific comments included liking the text reminders for the diary and to remember to wear the devices (1). Participants offered ways of improving the study including making the device smaller and addressing challenges with the headache diary only being once a day but experiencing multiple headache spikes a day.

## Discussion

We conducted a study measuring light exposure during everyday life of 20 youth with migraine, most with high migraine disease burden. To our knowledge, this is the first study demonstrating light exposure habits in a population with migraine using wearable light logger technology. The goal of this study was to demonstrate the feasibility of collecting light logger measurements and provide data to determine sample sizes needed for larger studies. Here, we review intriguing findings that are beginning to emerge in youth with migraine in the context of other studies, and how these findings should inform future study design.

We found that participants spent only about 15% of the daytime (1.5 hours per day) at or above the recommended minimum daylight levels. By comparison, they spent most of their time below the recommended maximum light levels 3 hours before bed, and during the night (78% and 99% on average, respectively). We suspect this is due to multiple factors. Perhaps one of the largest contributors to low daytime light exposure— not unique to individuals with photophobia—is the tendency in the modern societies to spend most of the time in indoor lighting environments.

A prior study by Lucas and colleagues provides an important reference point. They obtained continuous light exposure data from 59 healthy, mostly younger adults in Manchester, England. Similar to our study, they found participants spent a median of 3 of 9 daylight hours (33%) at or above recommended light levels, but 66% of the time at or below recommended light levels 3 hours before bed.^18^ Notably, however, participants in our study had still lower light exposure levels throughout the recording period (day, pre-bed, and nighttime). The healthy adults spent an average of 1.7 hours per day under bright light conditions (>1,000 lux mEDI),^18^ compared to an average of 40 minutes per day in our study. These differences are present even though Lucas and colleagues used wrist watches that can underestimate light exposure compared to pendant wear.

While it is possible that the low light exposure levels we recorded reflects light avoidance driven by migraine-related photophobia in our study population, there are other explanations. In addition to differences in age in the studied groups, Lucas and colleagues completed recording February through July, while we recorded November through March, biasing our study towards the winter months when light exposure tends to be lower. Indeed, a study including 15 adults in Amsterdam found a 4-fold difference between time spent above 250 lux mEDI between the winter and summer.^13^ We also found that outdoor light exposure was variable across participants, ranging from 0 to almost 2 hours a day. This may reflect differences in other factors like extracurricular activities (e.g. participation in outdoor sports) that could influence light exposure. Clearly, simultaneous measurements of study and control populations will be needed to support stronger claims of altered visual diet in migraine.

There is a strong relationship between light exposure and the sleep/wake cycle. Several features of our data are consistent with prior studies. We observed that youth had a later shift in light exposure profiles on the weekends compared to the weekdays. This corresponded with reported bedtime, sleep time, and wake-up times. Our findings are consistent with prior reports that find exposure to artificial light before bed is associated with sleep disruption, and reduced light exposure during the daytime is associated with daytime sleepiness.^18,33^ Additionally, mean melanopic illuminance in the 3 hours around “lights out” (9pm – 12am) was generally higher in youth with chronic migraine, moderate-to-severe headache-related disability, and high light sensitivity. Furthermore, the entire temporal light profile was shifted an hour later for youth with chronic migraine. Interestingly, later chronotype has been associated with more frequent headache in youth, consistent with these findings.^12^ The causal direction of these associations is unknown. It is possible, for example, that youth with chronic migraine arise later due to severe symptoms, producing a shifted sleep schedule, resulting in greater artificial light exposure in the evening and before bed. Alternatively, these effects may reflect sleep disruption influenced by evening and nighttime screen use. This is consistent with the finding that increased headache frequency in youth is associated with prolonged screen use,^11,12^ though these findings are based on self-report and further confirmation is needed.^34^

Interestingly, there was no relationship between self-reported fear-of-pain and sensory avoidant behavior and light exposure. This was despite the generally high reports of fear-of-pain and avoidance in this group, consistent with chronic pain conditions including migraine.^32^ Overall, reports of light avoidant behavior may not fully reflect actual daily habits, highlighting the need for objective measurements.

Power analyses revealed that sample sizes of 100 to 150 are needed to detect group differences in light metrics for migraine frequency and disability, light sensitivity, and sleep (assuming that the sub-group differences we observed in our pilot data are veridical). Prior reports have offered that significant differences between winter and summer light exposure should be found with sample sizes as small as 3 individuals.^13^ Our calculations in youth with migraine indicate differences within disease states may be more subtle, requiring larger sample sizes.

### Strengths and Limitations

To our knowledge, this is the first study to report objective measurements of light exposure combined with a daily diary to track migraine symptoms in real time. We used a light logger with 10 channels of differing spectral sensitivities allowing for the separation of photopic and melanopic illuminance, while light loggers with a single channel provide only a non-specific measure of overall illuminance. The multi-channel measurements allow for more mechanistic hypothesis testing. Sudden increases in both photopic and melanopic lux can contribute to visual discomfort.^35^ However, melanopic signals have a greater effect on circadian entrainment than photopic signals, while manipulation of photopic illuminance levels is the goal of artificial lighting.^26^ And while photopic and melanopic illuminance intensities are correlated in natural light, they can become relatively dissociated under artificial lighting.^36^ making it advantageous to measure both. We used melanopic signals to measure the timing of light exposure, including how this relates to sleep/wake habits, while we used photopic signals to measure the overall intensity of light exposure in a 24-hour period, and as a threshold to separate brightness typical of outdoor versus indoor light. Measurements were also collected as a pendant around the neck that improves the accuracy of light measurements.^14,24^ Overall, diary and device compliance were high providing a complete dataset for analysis, and participants generally had positive feedback on the study design.

Limitations include small sample size and the lack of a control group. Furthermore, participants were established patients of a pediatric headache clinic receiving active treatment, limiting the generalizability of the results. There may be bias in that youth willing to participate in the study may be more likely to already be working on healthy headache habits, while those still struggling with daily habits may be less likely to participate. While light filtering lens use was captured by survey data, we did not correct our measurements for the continuous use of blue light filtering glasses for one participant. Finally, data collection occurred in the winter months when light exposure is lower in the Philadelphia area thus it is unclear if results are generalizable to different times of year.

### Future Directions

This study demonstrates the feasibility of recording daily light exposure in youth with migraine. Furthermore, we found intriguing differences between youth with low compared to high frequency (chronic) migraine. Future studies that include larger sample sizes and control comparison to migraine-free peers are needed. Based on our pilot data, sample sizes of 100 – 150 participants would be sufficient for most light exposure metrics to identify group differences in youth with migraine. Collection of additional information including extracurricular activities and access to green spaces that could impact light exposure should also be included. Controlling for time of year is also critical as there are dramatic differences in light exposure between the summer and winter months at greater latitudes.^37^ This cycle may contribute to the seasonal variation observed in migraine, where migraine symptoms to be worse in the late fall and winter months,^38–41^ and seasonal comparison offers a unique opportunity for within-participant measurement.

Overall levels of sleep impairment and disturbance were high in our sample. While light exposure generally agreed with reported chronotype, it is important to note that individuals with migraine tend to overestimate how long it takes for them to fall asleep, indicating migraine may affect the perception of sleep.^42^ Combining continuous light logger technology with actigraphy that can measure sleep, and physical activity could provide detailed, reliable, and objective information about daily habits that could be used to develop circadian rhythm profiles. This approach has the potential to serve as the basis for clinical trials that use biosensor feedback to measure treatment response, and to develop evidence-based, personalized non-pharmacologic management approaches to healthy headache habits.

## Conclusion

Similar to prior studies in a general adult population, youth with migraine receive less than recommended daytime light exposure but do tend to achieve recommended levels of darkness during the evening and night. Associating migraine symptoms with daily light exposure in youth is feasible and demonstrates intriguing differences. Further study with larger sample sizes of 50 to 150, inclusion of timepoints across the school year when natural daylight hours are different, and comparison to migraine-free controls is needed.

## Conflict of interest statement

### Commercial Relationships Disclosures

C.P.G.: Dr. Patterson Gentile is currently funded by the National Institutes of Health/National Institute of Neurological Disorders and Stroke (K23 NS124986) and the CHOP Foerderer Institutional grant.

C.L.S.: Dr. Szperka has received research/grant support from the National Institutes of Health/National Institute of Neurological Disorders and Stroke (K23 NS102521), and PCORI. Dr. Szperka or her institution have received compensation for her consulting work for Eli Lilly; Teva Pharmaceutical Industries Ltd; Upsher-Smith Laboratories, LLC; and Abbvie.

A.D.H.: Dr. Hershey or his institution have received compensation for serving as a consultant for AbbVie, Amgen, Biohaven, Eli Lilly, Lundbeck, Supernus, Teva, Theranica and Upsher-Smith. His institution has also received research support from Amgen, Biohaven, Eli Lilly, Theranica, Upsher-Smith, and the NIH NINDS/NICHDS.

G.K.A.: Dr. Aguirre receives funding/grant support from the National Institute of Neurological Disorders and Stroke, the National Eye Institute, and the Binational Science Foundation.

R.S., B.M.P., and N.R. do not have conflicts of interest.

## Acknowledgements

We would like to thank the participants for contributing their time and feedback to this study.

## Abbreviations

CHOP: Children’s Hospital of Philadelphia
CHD-3: International Classification of Headache Disorders 3^rd^ Edition
IQR: interquartile range
mEDI: Melanopic Equivalent Daylight Illuminance
SD: standard deviation

## References

1. Digre KB, Brennan KC. Shedding light on photophobia. Journal of Neuro-Ophthalmology 2012.

2. Patterson Gentile C, Szperka CL, Hershey AD. Cluster Analysis of Migraine-associated Symptoms (CAMS) in youth: A retrospective cross-sectional multicenter study. Headache: The Journal of Head and Face Pain. 2024;64:1230–1243.

3. Rogers D, Protti T, Ngo B, et al. Interictal Avoidance Of Migraine-Related Stimuli. J Pain. 2023;24:48–49.

4. Windred DP, Burns AC, Lane JM, et al. Brighter nights and darker days predict higher mortality risk: A prospective analysis of personal light exposure in >88,000 individuals. Proceedings of the National Academy of Sciences. 2024;121.

5. Ruby NF, Brennan TJ, Xie X, et al. Role of Melanopsin in Circadian Responses to Light. Science (1979). 2002;298:2211–2213.

6. Lu J, Zou R, Yang Y, et al. Association between nocturnal light exposure and melatonin in humans: a meta-analysis. Environmental Science and Pollution Research. 2023;31:3425–3434.

7. Smolensky MH, Sackett-Lundeen LL, Portaluppi F. Nocturnal light pollution and underexposure to daytime sunlight: Complementary mechanisms of circadian disruption and related diseases. Chronobiol Int. 2015;32:1029–1048.

8. Mireku MO, Barker MM, Mutz J, et al. Night-time screen-based media device use and adolescents’ sleep and health-related quality of life. Environ Int. 2019;124:66–78.

9. Šmotek M, Fárková E, Manková D, Kopřivová J. Evening and night exposure to screens of media devices and its association with subjectively perceived sleep: Should “light hygiene” be given more attention? Sleep Health. 2020;6:498–505.

10. Vijakkhana N, Wilaisakditipakorn T, Ruedeekhajorn K, Pruksananonda C, Chonchaiya W. Evening media exposure reduces night-time sleep. Acta Paediatr. 2015;104:306–312.

11. Montagni I, Guichard E, Carpenet C, Tzourio C, Kurth T. Screen time exposure and reporting of headaches in young adults: A cross-sectional study. Cephalalgia. 2016;36:1020–1027.

12. Nilles C, Williams J V., Patten SB, Pringsheim TM, Orr SL. Lifestyle Factors Associated With Frequent Recurrent Headaches in Children and Adolescents. Neurology. 2024;102.

13. Zauner J, Udovicic L, Spitschan M. Power analysis for personal light exposure measurements and interventions. PLoS One. 2024;19:e0308768.

14. Spitschan M, Smolders K, Vandendriessche B, et al. Verification, analytical validation and clinical validation (V3) of wearable dosimeters and light loggers. Digit Health. 2022;8:205520762211448.

15. Guidolin C, Udovicic L, Broszio K, et al. Protocol for a prospective, multi-centric, cross-sectional 2 cohort study to assess personal light exposure. Medrxiv. Epub 2024.

16. Harris PA, Taylor R, Thielke R, Payne J, Gonzalez N, Conde JG. Research electronic data capture (REDCap)—A metadata-driven methodology and workflow process for providing translational research informatics support. J Biomed Inform [online serial]. 2009;42:377–381. Accessed at: https://linkinghub.elsevier.com/retrieve/pii/S1532046408001226.

17. Harris PA, Taylor R, Minor BL, et al. The REDCap consortium: Building an international community of software platform partners. J Biomed Inform [online serial]. 2019;95:103208. Accessed at: https://linkinghub.elsevier.com/retrieve/pii/S1532046419301261.

18. Didikoglu A, Mohammadian N, Johnson S, et al. Associations between light exposure and sleep timing and sleepiness while awake in a sample of UK adults in everyday life. Proceedings of the National Academy of Sciences. 2023;120.

19. Szperka CL, Farrar JT, Hershey AD. Improving headache diagnosis and treatment through patient headache questionnaires. American Headache Society Virtual Annual Scientific Meeting;

20. Verriotto JD, Gonzalez A, Aguilar MC, et al. New Methods for Quantification of Visual Photosensitivity Threshold and Symptoms. Transl Vis Sci Technol. 2017;6:18.

21. van Kooten JAMC, van Litsenburg RRL, Yoder WR, Kaspers GJL, Terwee CB. Validation of the PROMIS Sleep Disturbance and Sleep-Related Impairment item banks in Dutch adolescents. Quality of Life Research. 2018;27:1911–1920.

22. Werner H, LeBourgeois MK, Geiger A, Jenni OG. Assessment of Chronotype in Four- to Eleven-Year-Old Children: Reliability and Validity of the Children’s ChronoType Questionnaire (CCTQ). Chronobiol Int. 2009;26:992–1014.

23. Kellier DJ, Marquez de Prado B, Haagen D, et al. Development of a text message-based headache diary in adolescents and children. Cephalalgia. 2022;42:1013–1021.

24. Figueiro M, Hamner R, Bierman A, Rea M. Comparisons of three practical field devices used to measure personal light exposures and activity levels. Lighting Research & Technology. 2013;45:421–434.

25. Dautovich ND, Schreiber DR, Imel JL, et al. A systematic review of the amount and timing of light in association with objective and subjective sleep outcomes in community-dwelling adults. Sleep Health. 2019;5:31–48.

26. Michael PR, Johnston DE, Moreno W. A conversion guide: solar irradiance and lux illuminance. Journal of Measurements in Engineering. 2020;8:153–166.

27. Knoop M, Stefani O, Bueno B, et al. Daylight: What makes the difference? Lighting Research & Technology. 2020;52:423–442.

28. Brown TM, Brainard GC, Cajochen C, et al. Recommendations for daytime, evening, and nighttime indoor light exposure to best support physiology, sleep, and wakefulness in healthy adults. PLoS Biol. 2022;20:e3001571.

29. International Headache Society. The international classification of headache disorders, 3rd edition. Cephalalgia. 2018;38:1–211.

30. Hershey AD, Powers SW, Vockell A-LB, LeCates S, Kabbouche MA, Maynard MK. PedMIDAS. Neurology. Epub 2001.

31. Heyer GL, Perkins SQ, Rose SC, Aylward SC, Lee JM. Comparing patient and parent recall of 90-day and 30-day migraine disability using elements of the PedMIDAS and an Internet headache diary. Cephalalgia. 2014;34:298–306.

32. Simons LE, Sieberg CB, Carpino E, Logan D, Berde C. The Fear of Pain Questionnaire (FOPQ): Assessment of Pain-Related Fear Among Children and Adolescents With Chronic Pain. J Pain. 2011;12:677–686.

33. Burns AC, Saxena R, Vetter C, Phillips AJK, Lane JM, Cain SW. Time spent in outdoor light is associated with mood, sleep, and circadian rhythm-related outcomes: A cross-sectional and longitudinal study in over 400,000 UK Biobank participants. J Affect Disord. 2021;295:347–352.

34. Langdon RL, DiSabella MT, Strelzik JA. Screen time and pediatric headache: A scoping review of the literature. Headache: The Journal of Head and Face Pain. 2024;64:211–225.

35. McAdams H, Kaiser E, Igdalova A, Brainard DH, Aguirre GK. Migraine is associated with greater sensitivity to melanopsin and cone stimulation. J Vis. 2019;19:22.

36. Longcore T, Rodríguez A, Witherington B, Penniman JF, Herf L, Herf M. Rapid assessment of lamp spectrum to quantify ecological effects of light at night. J Exp Zool A Ecol Integr Physiol. 2018;329:511–521.

37. Thorne HC, Jones KH, Peters SP, Archer SN, Dijk D-J. Daily and Seasonal Variation in the Spectral Composition of Light Exposure in Humans. Chronobiol Int. 2009;26:854–866.

38. Shin Y, Park H, Shim J, Oh M, Kim M. Seasonal Variation, Cranial Autonomic Symptoms, and Functional Disability in Migraine: A Questionnaire-Based Study in Tertiary Care. Headache: The Journal of Head and Face Pain. 2015;55:1112–1123.

39. Pakalnis A, Heyer GL. Seasonal Variation in Emergency Department Visits Among Pediatric Headache Patients. Headache: The Journal of Head and Face Pain. 2016;56:1344–1347.

40. Caperell K, Pitetti R. Seasonal Variation of Presentation for Headache in a Pediatric Emergency Department. Pediatr Emerg Care. 2014;30:174–176.

41. Radziwon J, Waszak P. Seasonal changes of internet searching suggest circannual rhythmicity of primary headache disorders. Headache: The Journal of Head and Face Pain. 2022;62:811–817.

42. Rantanen O, Hollmen M, Bachour A. Migraine may disturb sleep perception during sleep onset: a retrospective data analysis. Journal of Clinical Sleep Medicine. 2022;18:2113–2117.

